# Acute inhibition of iron-sulfur cluster biosynthesis disrupts metabolic flexibility in mice

**DOI:** 10.1101/2024.08.19.608291

**Authors:** Marte Molenaars, Hannan Mir, Samantha W. Alvarez, Lakshmi Arivazhagan, Carolina Rosselot, Di Zhan, Christopher Y Park, Adolfo Garcia-Ocana, Ann Marie Schmidt, Richard Possemato

**Author notes:** Equal contribution (Marte Molenaars, Hannan Mir).

## Abstract

Iron-sulfur clusters (ISCs) are cell-essential cofactors present in ∼60 proteins including subunits of OXPHOS complexes I-III, DNA polymerases, and iron-sensing proteins. Dysfunctions in ISC biosynthesis are associated with anemias, neurodegenerative disorders, and metabolic diseases. To assess consequences of acute ISC inhibition in a whole body setting, we developed a mouse model in which key ISC biosynthetic enzyme NFS1 can be acutely and reversibly suppressed. Contrary to in vitro ISC inhibition and pharmacological OXPHOS suppression, global NFS1 inhibition rapidly enhances lipid utilization and decreases adiposity without affecting caloric intake and physical activity. ISC proteins decrease, including key proteins involved in OXPHOS (SDHB), lipoic acid synthesis (LIAS), and insulin mRNA processing (CDKAL1), causing acute metabolic inflexibility. Age-related metabolic changes decelerate loss of adiposity substantially prolonged survival of mice with NFS1 inhibition. Thus, the observation that ISC metabolism impacts organismal fuel choice will aid in understanding the mechanisms underlying ISC diseases with increased risk for diabetes.

**Graphical abstract:** 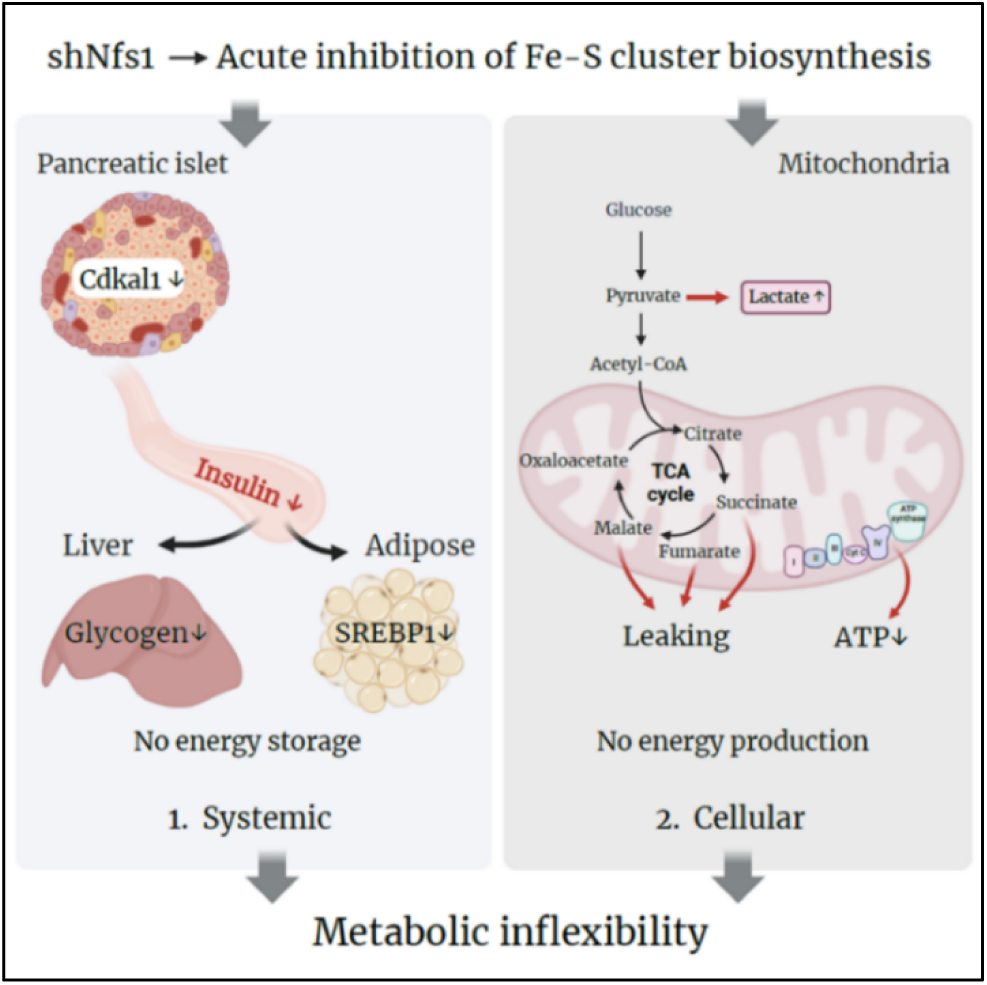

**Highlights:**

- Acute ISC inhibition leads to rapid loss of adiposity in mice
- Multi-metabolic pathway disruption upon ISC deficiency blocks energy storage
- Nfs1 inhibition induces glucose dyshomeostasis due to ISC deficiency in β-cells
- Energy distress caused by inhibition of ISC synthesis is attenuated in aged mice

## Introduction

Among the myriad functions of the mitochondria, synthesis of iron-sulfur cluster (ISC) cofactors has been considered the most irreplaceable.^1,2^ ISCs are cell-essential cofactors present in at least 60 proteins and are required for their diverse functions, including as electron carriers in the electron transport chain (ETC), DNA replication, mRNA transcription, and iron sensing.^3^ Eukaryotic ISC biosynthesis begins with the removal of sulfur from cysteine by the enzyme Nitrogen Fixation 1 Cysteine Sulfurase (NFS1) and its combination with iron on scaffold protein ISCU. Subsequently, numerous chaperones and maturation factors are responsible for the generation of [4Fe-4S], [3Fe-4S], and [2Fe-2S] clusters and their delivery to ISC-proteins.^3,4^ These ISC-proteins include ETC Complexes I, II, and III, the catalytic subunits of the replicative DNA polymerases (POLA, POLD, and POLE), several helicases involved in DNA repair, mitoribosomal assembly factor METTL17, the TCA enzyme aconitase (ACO2), heme synthesis enzyme FECH, and lipoic acid synthesis enzyme LIAS. Besides being important for individual ISC proteins, the availability of ISCs also plays a role in iron sensing and regulation of iron levels via iron regulatory proteins 1 and 2 (IRP1 and IRP2), albeit by different mechanisms. IRP1, encoded by the gene ACO1, functions as a cytoplasmic aconitase when bound to an ISC,^5^ and loss of this ISC reveals an RNA binding domain that recognizes hairpin-like structures called iron-responsive elements (IRE).^6^ In contrast, IRP2 stability and IRE-binding activity is inhibited by ISCs.^7,8^ IRP targets are mainly involved in iron metabolism and include transferrin receptor (TFRC) and ferritin subunits FTH1 and FTL, regulating iron import and storage respectively.^6,9–14^ Thus, ISCs are intimately linked to the cellular iron sensing apparatus, as reduced ISC abundance enhances iron availability by regulating iron transport and storage, which we term the iron starvation response.

Dysfunctions in ISC biosynthesis are associated with multiple human diseases whose presentation varies depending on the perturbation. Rare loss-of-function mutations in NFS1 cause perinatal mitochondrial disease^15^ and mutations in ISCU occur in patients exhibiting hereditary myopathy with lactic acidosis.^16^ Mutations in genes selectively required for mitochondrial ISC maturation underlie multiple mitochondrial dysfunctions syndrome, including NFU1, BOLA3, IBA57, ISCA1, and ISCA2.^16^ Sideroblastic anemia, characterized by the production of iron-loaded erythroid precursors (ringed sideroblasts), can be caused by mutations in genes directly impacting heme biosynthesis (e.g. ALAS2 and SLC25A38), which requires the ISC protein ferrochelatase (FECH), or by mutation of genes selectively impacting cytosolic ISC biogenesis, including GLRX5, HSPA9, and ABCB7.^17^ FXN is mutated in Friedreich’s Ataxia, the most common monogenic mitochondrial disorder, and in contrast to the above phenotypes, is characterized by progressive gait and limb ataxia that develops over decades.^18^ The exact function of FXN is debated, and it has been described as both an iron oxidoreductase and an allosteric activator of NFS1.^19–21^ Friedreich’s Ataxia patients are at a 3-fold increased risk for developing diabetes, however it is unclear how this effect is related to ISC deficiency.^22^

Altogether, the clinical consequences for patients with loss of ISC biosynthetic proteins can vary widely, either because of the precise ISC containing proteins affected, the way in which ISC biogenesis is affected, or because ISC formation and stability are highly dependent on the cell type and local environment, particularly environmental oxygen concentration. Molecular oxygen disrupts ISCs through oxidative damage, leading to the breakdown of the cluster. As such, the effects of ISC inhibition are dependent on environmental oxygen concentration, both in cells and tissues.^23,24^ For example, acute inhibition of ISC biosynthesis in HEK293 cells rapidly increases citrate levels due to loss of ISC protein aconitase (ACO2), leading to increased fatty acid biosynthesis and lipid droplet accumulation.^25^ However, as many *in vitro* experiments studying ISC biosynthesis are carried out under atmospheric oxygen levels (21%), these results might not reflect *in vivo* mechanisms of ISC formation and stability where most cells experience much lower oxygen tension (∼5%) and have tissue-specific metabolic demands. For these reasons, we developed a mouse model in which NFS1 can be acutely suppressed in adult animals in order to better assess consequences of ISC inhibition in a whole body holistic setting, which might better reflect ISC related pathologies.

Here, we find that acute inhibition of iron-sulfur biosynthesis *in vivo* leads to activation of cell-type specific mitochondrial stress responses and rapid metabolic rewiring resulting in loss of white adipose tissue (WAT). Upon NFS1 knockdown (KD), both female and male mice exhibit a metabolic shift toward lipid catabolism and liver glycogen depletion without changes in physical activity and caloric intake. While other mouse models with mitochondrial disturbances shift to using glucose as a fuel source, NFS1 KD results in insulin insufficiency and glucose intolerance. ISC protein CDKAL1, involved in insulin processing, is downregulated in isolated pancreatic islets upon NFS1 KD, as are SDHB and LIAS, important for TCA cycle and OXPHOS function. Finally, we show that (advanced) aging rescues survival of mice upon NFS1 KD; in contrast, simply increasing caloric intake and adiposity do not result in rescue. These observations demonstrate that ISC metabolism contributes to organismal fuel choice by regulating insulin production and mitochondrial metabolism.

## Results

### Acute inhibition of ISC synthesis causes weight loss, tissue-specific stress responses, and suppression of lipid biosynthesis

In order to investigate the effect of *in vivo* inhibition of ISC biosynthesis, we developed a mouse model in which the key ISC synthesis enzyme NFS1 can be suppressed, using targeted alleles in which a tetracycline responsive promoter drives a miR30-based, validated Nfs1 shRNA (Fig 1A).^26^ Administration of doxycycline (50 μg/ml in drinking water) to 20-week-old mice reduced NFS1 protein levels in multiple tissues (Fig 1B). After two weeks, mice lost around 20% of their initial weight, requiring euthanasia (Fig 1C-D), in line with the premise that sufficient ISC biosynthesis is indispensable in organisms.

**Figure 1.**
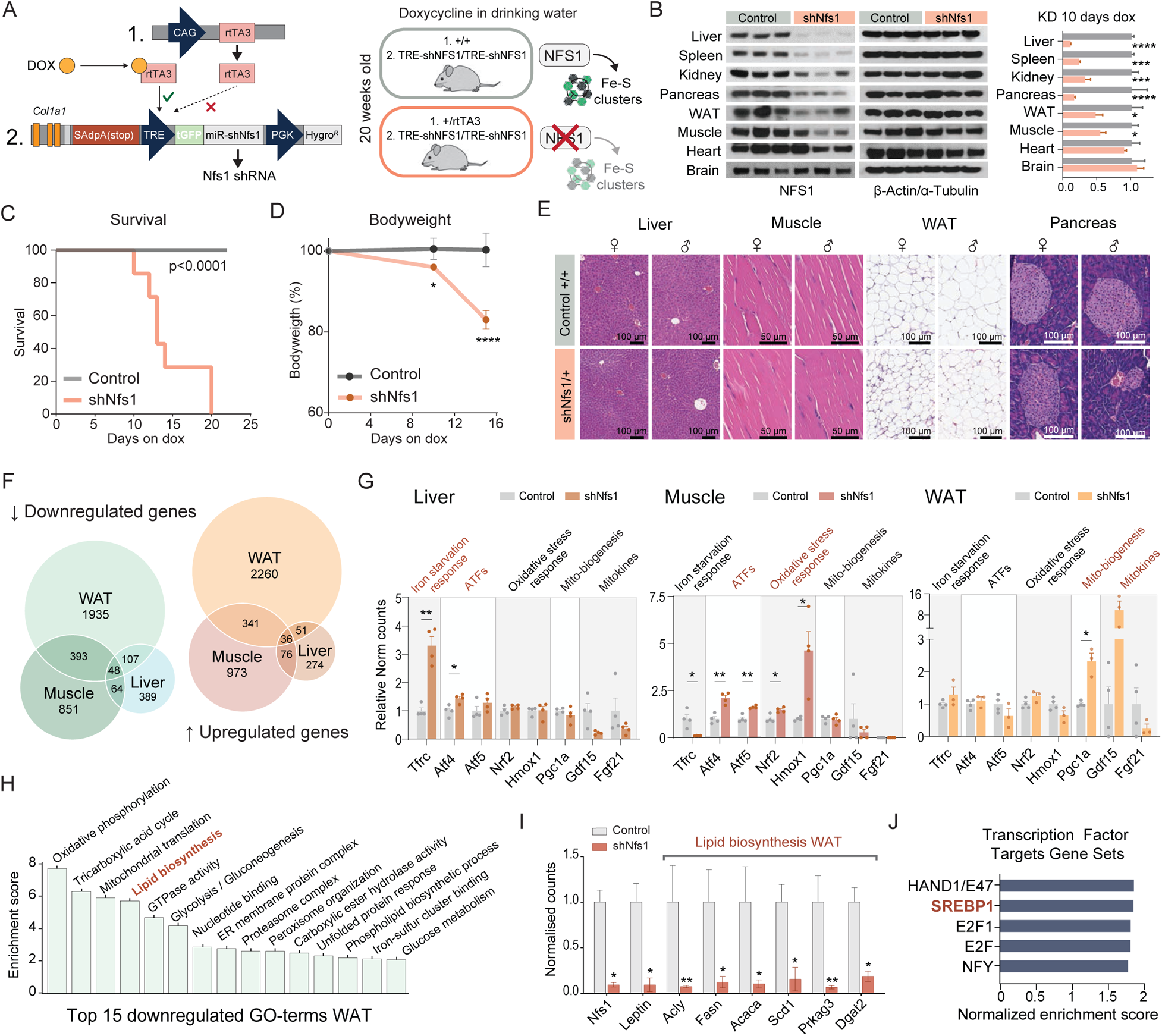
Acute inhibition of ISC synthesis causes weight loss, tissue-specific stress responses, and suppression of lipid biosynthesis. A) Schematic of the Nfs1 knockdown mouse model. B) Immunoblot of NFS1 and B-actin after 10 days of doxycycline (dox) treatment in indicated tissues, n=3 for both groups. C) Survival curve of 20-week-old control versus Nfs1-suppressed mice, n=10 for both groups. D) Bodyweight % of 20-week-old control versus Nfs1-suppressed mice, n=4 for both groups. E) H&E-stained sections indicated tissues of control and Nfs1-suppressed mice after 10 days of dox addition, female (♀) and male (♂). F) Venn diagram of differentially expressed mRNAs in WAT (1935↓ and 2260↑), Muscle (851↓ and 973↑) and Liver (389↓ and 274↑) after 10 days of dox addition, measured using RNAseq. G) Expression of mRNAs related to the indicated stress responses in Liver, Muscle, and eWAT, measured using RNAseq, n=3 for eWAT Nfs1-suppressed mice and n = 4 for all other groups. H) The top 15 enriched GO-terms downregulated in WAT using DAVID analysis clusters enrichments in Gene Ontologies (GO). I) Expression of Nfs1, Leptin, and indicated lipid biosynthesis genes in eWAT of control vs Nfs1-suppressed mice, measured using RNAseq with n=3 for Nfs1-suppressed mice and n=4 for control group. J) Enrichment scores for transcription factors associated to transcriptional changes in WAT of control vs Nfs1-suppressed mice. Data are presented as mean ± SEM, comparisons made using two-way analysis of ANOVA (B, D, J) or Log-rank (Mantel-Cox) test (C) or multiple unpaired t tests (I); *p < 0.05, **p < 0.01, ***p < 0.001, ****p < 0.0001.

As heme and hemoglobin synthesis are dependent on ISC biogenesis and one of the pathologies seen in patients with defective ISC biosynthesis is anemia, we hypothesized that NFS1 KD could similarly disrupt hematopoietic cell function. However, in Nfs1-suppressed mice we did not observe differences in red blood cell (RBC), basophil, eosinophil, or monocyte counts, hemoglobin, RBC distribution, mean cell hemoglobin concentration, or mean RBC volume, but a progressive increase in the ratio of total neutrophils to lymphocytes was observed over time (Fig S1A-B). A high neutrophil-to-lymphocyte ratio (NLR) indicates general physiological stress, and has been described as a biomarker for poor prognosis in many different types of disease.^27^

These data suggest that rather than disrupted erythropoiesis or anemia, other disturbed physiological processes are more predominant in these mice. Indeed, a bone marrow transplant (BMT) from wild type donor mice into lethally-irradiated Nfs1-suppressed mice did not rescue the weight loss nor the increased NLR (Fig S1C), demonstrating that defects in erythropoiesis or immune cells function do not underlie these phenotypes. Moreover, as neutrophils have a short half-life requiring their continuous production, these results indicate that the level of NFS1 suppression in this compartment is not sufficient to limit the cell proliferation and differentiation processes required to produce mature neutrophils.

To identify other potential causes for the physiological stress and weight loss in Nfs1-suppressed mice, we analyzed histology of liver, muscle, WAT, and pancreas using hematoxylin and eosin (H&E) stain after 10 days of NFS1 KD. The architecture of these tissues was not observably altered, with the exception of pancreatic islet β-cells, which exhibited a reduced cytoplasmic density (Fig 1E, Fig S1D). As insulin represents approximately 10% of total β-cell protein content, reduced cytoplasmic density could indicate reduced insulin production.^28^

To understand which biological processes might be altered upon acute inhibition of ISC biosynthesis, we performed RNA sequencing (RNA-seq) analysis in liver, epididymal white adipose tissue (eWAT), and skeletal muscle isolated from control and Nfs1-suppressed mice 10 days after initiation of DOX (Fig S1E, Table S1). We observed substantial gene expression changes in these tissues, with the greatest number of significant changes in eWAT with 1935 mRNAs downregulated and 2260 mRNAs upregulated (Fig. 1F). Using DAVID analysis to cluster Gene Ontology (GO) enrichments, we observed that GO-terms related to transcription regulation and DNA damage are upregulated in multiple tissues (Fig S1H). This observation is in accordance with ISCs being a cofactor for transcription factor IIH (TFIIH) complex subunit ERCC2, Elongator complex member ELP3, replicative polymerases POLA, POLD, and POLE, and several DNA damage helicases, and with previously described *in vitro* work showing inhibition of either NFS1 or these polymerases in turn activates DNA damage responses.^29^ Collectively downregulated GO-terms in all tissues were related to mitochondrial function, consistent with the key roles of ISCs in the ETC and TCA cycle (Fig 1H, Fig S1F-G). Indeed, inhibiting ISC biogenesis affects mitochondrial function in cultured cells and is in line with perinatal mitochondrial phenotypes observed in patients with Nfs1 mutations.^15,30,31^

Processes specifically downregulated in the liver upon NFS1 inhibition included *Iron*, *Glutathione metabolic process* and *Insulin-activated receptor activity* (Fig S1G), including mRNAs related to iron and ISC metabolism such as *Ftl1*, *Ciao1*, *Ireb2*, and *Fdxn* (Table S1). We observed a parallel liver-specific upregulation of Tfrc, which is the predominant transcript containing an iron response element (IRE) in the 3’ UTR, indicating activation of the iron starvation response exclusively in the liver, known to be the site of systemic iron regulation (Fig 1G). We also observed activation of other stress responses in individual tissues, such as the integrated stress response (ISR) (Atf4, Atf5) in liver and muscle, oxidative stress (Nrf2, Hmox1) in muscle, and mitochondrial biogenesis (Pgc1a) and the mitokine Gdf15 in eWAT (Fig 1G). These stress responses are typically activated in models with reduced mitochondrial function and mitochondrial disease.^32–35^ However, Fgf21, a strong effector commonly activated by mitochondrial stress responses, was surprisingly unchanged (Fig 1F). Taken together, these data indicate that activation of stress responses by ISC inhibition are tissue and context-specific.

Interestingly, NFS1 inhibition induced the most number of significant transcriptional changes in eWAT, and one of the specifically downregulated GO-term was Lipid biosynthesis (Fig 2F). Downregulated mRNAs that are known to play a crucial role in lipid storage in adipose tissue included ATP citrate lyase (*Acly*), fatty acid synthase (*Fasn*), acetyl-CoA carboxylase (*Acaca*), stearoyl-Coenzyme A desaturase 1 (*Scd1*), AMP-activated protein kinase (*Prkag3*), and diacylglycerol O-acyltransferase 2 (*Dgat2*) (Fig. 2I). In addition, we observed a strong downregulation of the adipokine Leptin, an appetite suppressant that can be regulated by insulin, glucocorticoids, and cytokines (Fig. 2I). Transcription factor enrichment analysis (TFEA) revealed that SREBP1 targets are significantly enriched upon Nfs1 inhibition (Fig 1H). SREBP1, an insulin-mediated transcription factor, sustains adipocyte lipogenesis by facilitating expression of mRNAs such as *Acly*, *Acaca*, *Scd1*, and *Fasn.* Therefore, acute Nfs1 inhibition rapidly activates mitochondrial stress and iron starvation responses in a tissue-specific manner, and suppresses WAT lipid biogenesis.

**Figure 2.**
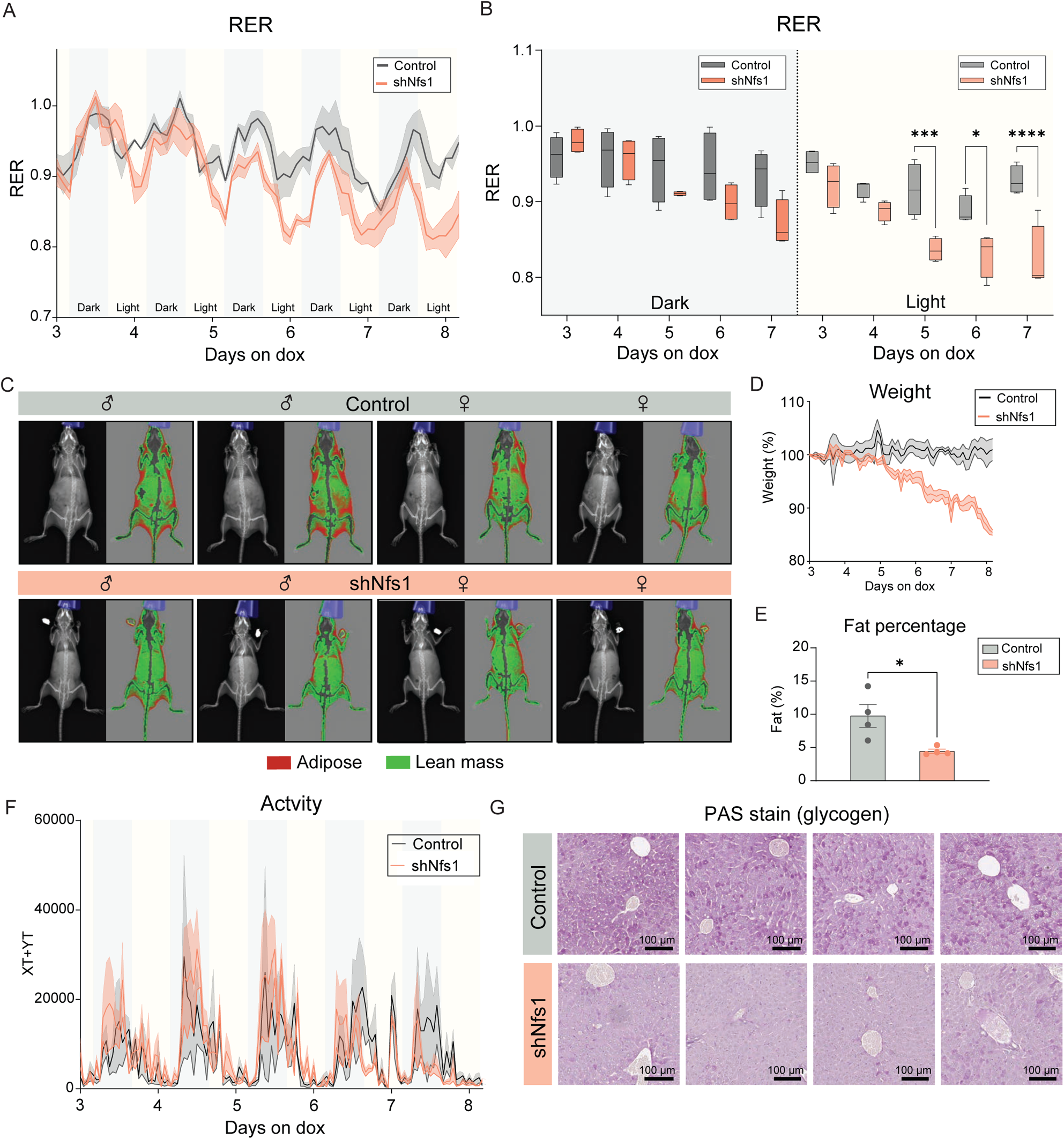
Respiratory exchange rate upon inhibition of ISC synthesis shifts towards lipid catabolism and is followed by body mass and fat loss. A) Respiratory exchange ratio (RER, VCO2/VO2) measured over a 5 day period between 3 and 8 days of dox. B) Average respiratory exchange ratio (RER, VCO2/VO2) during either dark or light period per day. C) Adipose and lean mass analyzed by dual energy X-ray absorptiometry (DEXA) scan. D) Body weight (%) was recorded between 3 and 8 days of dox. E) Body fat (%) measurements by DEXA. F) Physical activity was recorded between 3 and 8 days of dox. G) Glucose stored as glycogen measured by Periodic acid-Schiff reagent (PAS) in liver of control and Nfs1-suppressed mice after 10 days of dox. Data are presented as mean ± SEM; comparisons were made using two-way analysis of ANOVA, *p < 0.05, ***p < 0.001, ****p < 0.0001, n=4 for both groups

### Reduced respiratory exchange rate and fat percentage indicates increased lipid oxidation upon inhibition of ISC synthesis

To understand the metabolic consequences of ISC inhibition in vivo, we placed singly-housed mice in metabolic cages for analysis of O2 consumption, CO2 production, physical activity, and body weight. This analysis allowed us to calculate the respiratory exchange rate (RER), which reflects the ratio of carbon dioxide produced to oxygen consumed and thereby substrate utilization.

The initial RER values in both groups indicated an expected preference for carbohydrate utilization as the primary energy source (RER: 0.9-1) (Fig 2A). Beginning at 4 days after doxycycline addition, we observed a gradual decline in RER, demonstrating a shift towards increased lipid oxidation (Fig 2A). This effect was especially evident during the inactive light phase, and after 7 days the light cycle RER decreased to 0.8 (67% energy expenditure from fat oxidation) compared to 0.92 in the control condition (26% energy expenditure from fat oxidation, Fig 2B). Accordingly, dual energy X-ray absorptiometry (DEXA) scans show loss of adipose tissue upon Nfs1 inhibition (Fig. 2C). Indeed, NFS1 inhibition triggered a 15% loss of total body weight and >60% loss of body fat (Fig 2D-E), despite no observable changes in physical activity (Fig 2F), caloric intake, or core body temperature (Fig S2A-B). This decrease in adiposity was unanticipated as *in vitro* studies have shown that acute loss of ISCs results in increased fatty acid biosynthesis and generation of lipid droplets, indicating that systemic effects elicited by ISC deficiency are more critical than cell-intrinsic effects on metabolism in driving organismal metabolic changes to lipid biosynthesis.^25^ Moreover, histopathology revealed loss of liver glycogen stores upon NFS1 inhibition (Fig. 1G), but maintenance of muscle glycogen (Fig. S2C). This depletion could be caused by either impaired glycogen synthesis and/or storage mechanisms, as liver glycogen is synthesized upon increased glucose availability and insulin-mediated activation of glycogen synthesis.

As both long-term energy storage (fat) as well as short-term energy storage (glycogen) are not adequately maintained, we hypothesized that Nfs1 inhibition rapidly induces a metabolic shift downstream of alterations in nutrient sensing and fuel choice.

### Plasma metabolomics shows increased lactate and TCA intermediates and a dampened response to refeeding upon inhibition of ISC synthesis

ISCs are cofactors for multiple metabolic enzymes including OXPHOS complex I-III subunits and ACO2 (conversion of citrate to isocitrate in the TCA cycle) as well as the enzyme lipoate acid synthase (LIAS) required for the synthesis of lipoic acid, a cofactor for the pyruvate, branched-chain amino acid, glycine, and alpha-ketoglutarate dehydrogenase complexes. Cellular metabolic perturbations can result in altered uptake or secretion of metabolites and lipids, which can be identified using plasma metabolomics and lipidomics. Therefore, we measured the impact of Nfs1 KD on plasma metabolite and lipid levels from overnight fasted animals, as well as animals refed for 5 hours.

The top 25 most changed metabolites upon Nfs1 KD in fasted animals, ranked by partial least squares-discriminant analysis (PLS-DA) Variable Importance in Projection (VIP) score, included intermediates of the TCA cycle and lactate (Fig. 3A, Table S2). We also observed an increase in lactate in the refed state (Fig. 3B). We observed increased plasma levels of succinate, fumarate, and malate, especially in the refed state (Fig. 3B). However, we did not observe increased levels of citrate or isocitrate that were previously reported upon ISC inhibition *in vitro* attributed to inhibition of ISC protein ACO2.^25^ Elevated efflux of citrate to the cytoplasm can be used for fatty acid and lipid synthesis. However, we observed downregulation of lipid desaturase SCD1 in WAT (Fig. 1G), liver, and muscle (Table S2). While lipidomics analysis showed a significant increase in plasma triglycerides (TG) upon refeeding in control mice, Nfs1 inhibition did not significantly affect plasma TG levels compared to control mice in either the fasted or fed state (Fig. S3A-C, Table S3). Similarly, while FFAs were trending toward being decreased upon refeeding, Nfs1 inhibition did not significantly affect FFA levels in either the fasted or refed state (Fig. S3B-C, Table S3). Taken together, these data further demonstrate that TCA cycle metabolism and fatty acid synthesis alterations induced by ISC restriction in mice greatly diverge from the alterations observed in standard culture conditions.

**Figure 3.**
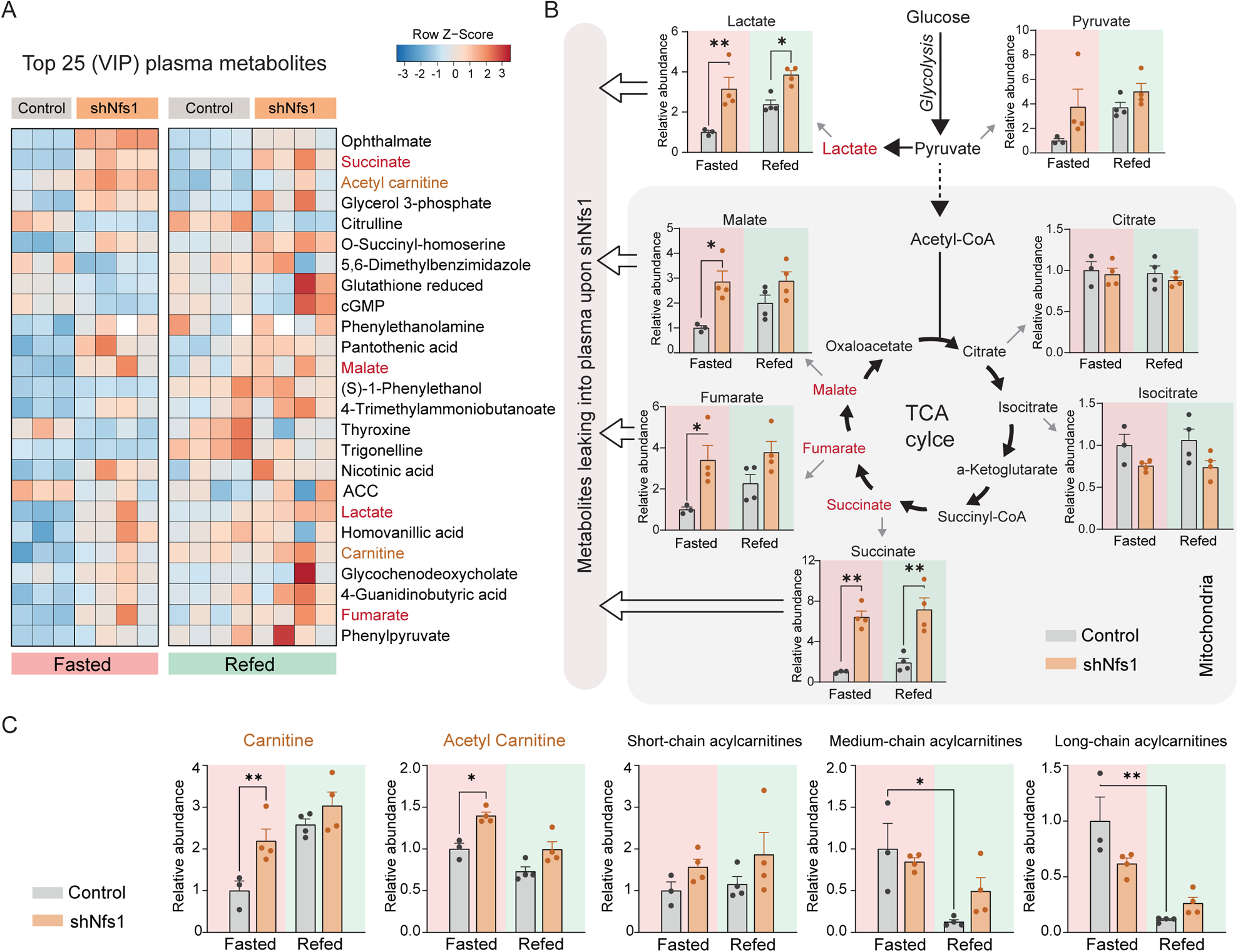
Plasma metabolomics shows increased lactate and TCA intermediates and a dampened response to refeeding upon inhibition of ISC synthesis. A) Heat map of most significantly changed metabolites in fasted state upon 10 days of dox treatment, ranked on VIP score. Fasted plasma on the left and refed plasma on the right, TCA-cycle intermediates are highlighted in red and carnitines are highlighted in orange. B) Individual metabolites of the TCA cycle during fasted and fed state and upon Nfs1 inhibition. C) Carnitine (LC), acetyl-Carnitine (C2), short-chain (C3-C5), medium-chain (C6-C12), long-chain (C14–C20) carnitines during fasted and fed state and upon Nfs1 inhibition. Data are presented as mean ± SEM; comparisons were made using two-way analysis of ANOVA, *p < 0.05, **p < 0.01, n=3 for fasted control mice and n=4 for all other groups.

Upon NFS1 inhibition, we also observed that carnitine and acetyl carnitine (C2) were significantly increased in the fasted state (Fig 3A). Moreover, upon refeeding, control mice exhibited decreased medium and long-chain acyl carnitine levels, an effect that was blunted upon NFS1 inhibition. Increased production of C2 represents a critical mechanism for buffering the metabolic status between fed (glucose oxidation) and fasted (fat oxidation) states, referred to as metabolic flexibility.^36^ It has been reported that persistent elevations in blood concentrations of C2 over time, as we observe upon NFS1 inhibition, may indicate systemic metabolic inflexibility.^37^

### Inhibition of ISC synthesis disturbs glucose homeostasis and insulin insufficiency contributing to an energy crisis that is attenuated by aging but not by high fat feeding

Metabolic flexibility, the ability of an organism to adjust to changes in the availability of energy substrates, is facilitated in part by the action of insulin. Because we observed morphologic changes to pancreatic islets (Fig. 1E) and gene expression changes in WAT consistent with reduced insulin signaling (Fig. 1I-J), we considered the extent to which glucose sensing by insulin is affected by inhibition of ISC synthesis.

Consistent with abnormal insulin function, upon NFS1 inhibition we observed elevated fasting blood glucose levels (Fig. 4A) and decreased plasma insulin levels, while glucose-dependent insulinotropic polypeptide (GIP) was increased (Fig. 4B). While GIP is released after meal ingestion to potentiate glucose-stimulated insulin release, the increased GIP upon NFS1 inhibition seems to be unable to elevate insulin levels.

**Figure 4.**
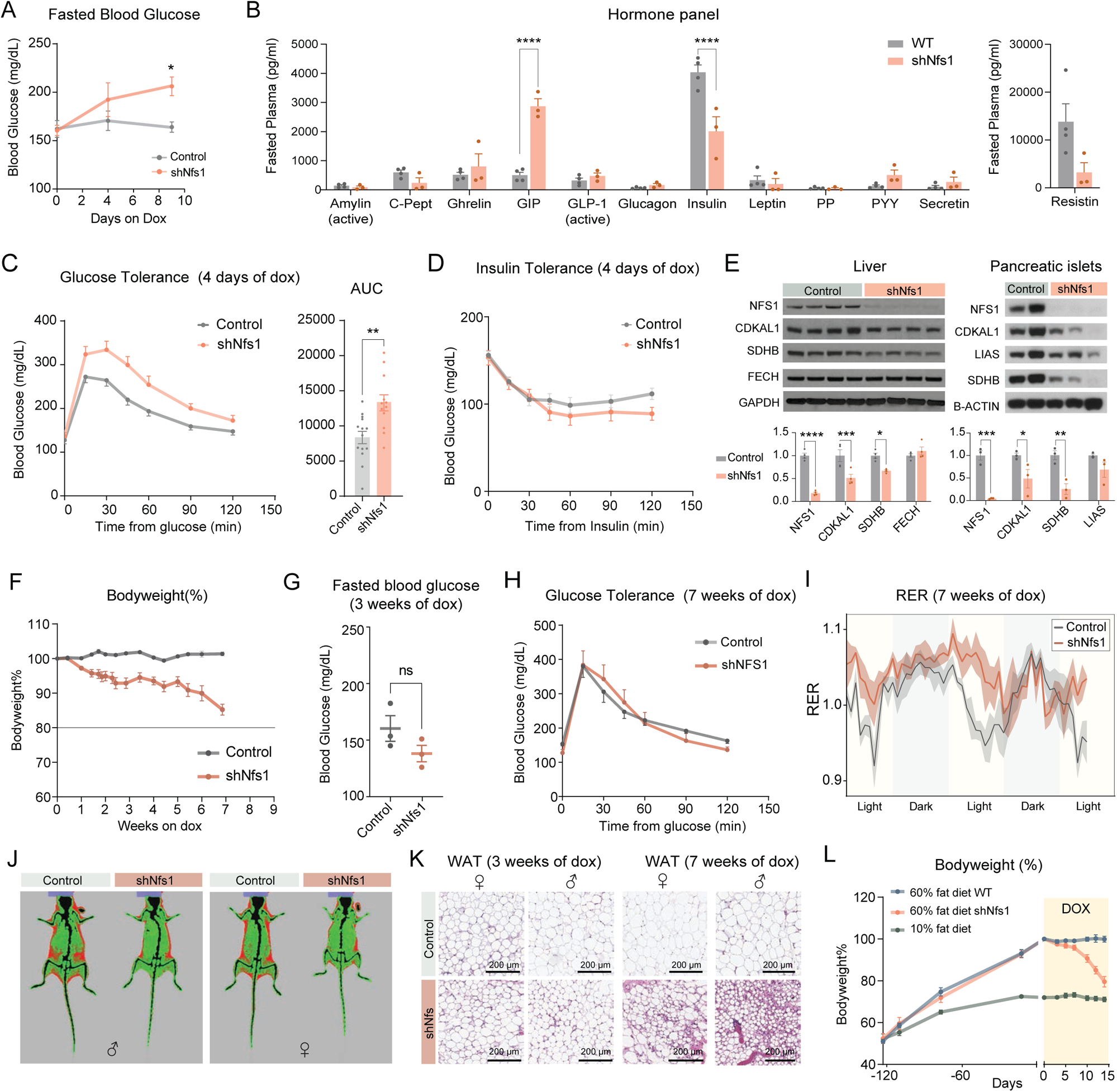
Inhibition of ISC synthesis disturbs glucose homeostasis and insulin insufficiency contributing to an energy crisis that is attenuated by aging but not by high fat feeding. A) Fasting glucose levels (6 hour fast) at indicated times after dox addition, n=4 for both groups. B) Metabolic hormone panel from 4 hour fasted plasma, with n=3 for Nfs1-suppressed mice and n=4 for control group. C) Oral Glucose tolerance test (OGTT) for Nfs1 inhibited and control mice after 4 days of dox. Blood glucose concentrations were determined at indicated times. Area under the curves shown at right, with n=12 for Nfs1-suppressed mice and n=14 for control group. D) Insulin tolerance test (ITT) for shNfs1 and control mice after 4 days of dox. Blood glucose concentrations were determined at indicated times. n=7 for both groups. E) Immunoblot of NFS1, ISC proteins (CDKAL1, SDHB, LIAS) in extracts of livers and isolated pancreatic islets from control and Nfs1-inhibited mice. Quantitation of protein levels normalized to loading control shown below, with n=4 for livers and n=3 for islets. F) Bodyweight (%) aged 1-year-old mice from Nfs1 inhibited and control mice, with n=10 per group. G) Fasting glucose levels (6 hour fast) in aged 1-year-old mice 3 weeks after dox addition. N=3 per group. H) Oral Glucose tolerance test (OGTT) for Nfs1 inhibited and control mice after 7 weeks dox treatment in 1-year old mice. n=7 for Nfs1-suppressed mice and n=8 for control group. I) Respiratory exchange ratio (RER, VCO2/VO2) in 1-year-old mice after 7 weeks of dox measured over ∼2 day period between with n=4 per group. J) Adipose and lean mass analyzed by dual energy X-ray absorptiometry (DEXA) scan from 1-year-old mice after 7 weeks of dox, female (♀) and male (♂). K) H&E-stained sections of eWAT from aged 1-year-old control and Nfs1-inhibited mice after 3 and 7 weeks of KD, female (♀) and male (♂). L) Bodyweight (%) control and Nfs1-inhibited male mice fed with 60% fat diet or fed with 10% fat control diet, with n=6 per group. Data are presented as mean ± SEM; comparisons were made using two-way analysis of ANOVA, *p < 0.05, **p < 0.01.

Accordingly, 4 days after induction of Nfs1 KD, mice show reduced glucose tolerance (Fig. 4C). In contrast, insulin stimulated glucose uptake by tissues is not affected, consistent with a primary insulin production defect (Fig. 4D). We therefore considered the degree to which pancreatic β-cell dysfunction and abnormal insulin signaling underlies the phenotypes observed upon Nfs1 inhibition. We observed that Nfs1 KD induced loss of SDHB, LIAS and CDKAL1 in isolated pancreatic islets (Fig 4E). Importantly, CDKAL1 is a type 2 diabetes susceptibility gene that catalyzes a tRNA modification important for insulin production. β-cell specific deletion of Cdkal1 reduces circulating glucose levels and decreases glucose tolerance similar to the phenotypes observed upon Nfs1 KD, but without loss of adiposity or body mass.^38^

If the phenotypes of ISC deficiency are due to signaling defects, as our model suggests, as opposed to toxicity in ß-cells or other tissues, we would expect that these phenotypes would be reversible following restoration of Nfs1 expression. Therefore, we treated mice with doxycycline for either 3, 6, or 9 days followed by withdrawal (Fig. S4A). All mice fully recovered their body weight upon doxycycline withdrawal, although a subset of mice treated with doxycycline for 9 days reached 20% body weight loss, requiring euthanasia. Upon 12 days of doxycycline withdrawal following 6 days of doxycycline treatment, we observed no abnormalities in OGTT (Fig. S4B). The reduced RER after 6 days of doxycycline upon Nfs1 inhibition normalized after ∼4 days of doxycycline withdrawal (Fig. S4D). We also did not observe any permanent tissue morphology changes, including changes to the cytoplasmic density in pancreatic islets (Fig. S4D). Therefore, the effects of NFS1 inhibition in this model are fully reversible, indicating a transient signaling defect.

As mice older than 10 months are known to have increased fat pad weight and doubled body fat percentage, as well as hyperinsulinemia related to increased insulin secretion,^39^ we hypothesized that the reversible metabolic inflexibility induced by ISC insufficiency could have distinct effects in older mice. Indeed, upon Nfs1 KD in 1-year-old mice, body mass loss was attenuated compared to 3-month old animals (Fig. 4F), despite similar KD efficiency (Fig S4E). Upon Nfs1 inhibition in 1-year-old mice we observed neither increased fasting glucose (Fig. 4G), nor morphological changes in pancreatic islets (Fig. S4E). While aged mice are known to have decreased glucose tolerance compared to young mice, we did not observe an additional effect of Nfs1 inhibition on glucose tolerance, even after 7 weeks of doxycycline treatment in older mice (Fig. 4H). Accordingly, the observed hormone changes in GIP and insulin upon NFS1 inhibition in young mice were blunted in aged mice (Fig. S4F). Moreover, RER was unaltered (Fig 4I), similar to other mouse models that exhibit decreased mitochondrial function without insulin insufficiency.^32^ Consistent with the attenuated loss of body mass, DEXA scans from 1-year-old mice show reduced adiposity after 7 weeks of doxycycline treatment (Fig. 4J). Interestingly, while we did not observe any morphological changes in the WAT of young mice nor in 1-year old mice after 3 weeks of doxycycline, we did observe decreased adipocyte size upon Nfs1 knockdown after 7 weeks of doxycycline (Fig 4K). These data indicate that the long-term effects of NFS1 inhibition on WAT in 1-year old mice, which may be related to reduced mitochondrial function, are divergent from the acute effects of NFS1 inhibition on WAT in young mice, which may be primarily driven by changes to insulin signaling.

Finally, we considered whether older mice might be protected from Nfs1 inhibition simply due to their relatively elevated body mass. High fat diet (HFD) feeding is a commonly used approach to induce rapid weight gain and, at extended time points, cause insulin resistance and diet-induced obesity. We placed mice on a high fat diet (60% kcal from lipids) or matched control diets (CD, 10% kcal from lipids) for 4 months, resulting in a 75% gain in body mass in the HFD fed mice compared to a 25% gain in body mass for the CD mice over that period (Fig. 4L). Inhibition of Nfs1 resulted in a similar rapid loss of body mass and adiposity in HFD mice, achieving humane endpoints at the same rate (Fig. 4L). Therefore, increased body mass and adiposity alone do not protect from ISC synthesis inhibition.

Altogether, we find that acute inhibition of ISC biosynthesis in adult mice leads to a reversible ‘double-hit’ on physiological metabolism, due to 1) reduced mitochondrial function and 2) reduced insulin secretion. ISC insufficiency induces canonical mitochondrial stress responses (with the exception of FGF21 production), and an iron-starvation response specifically in the liver. ISC insufficiency in pancreatic β-cells limits insulin secretion, suppressing eWAT fatty acid synthesis and liver glycogen storage and preventing a shift to glucose as the main energy source. Aged mice exhibit relative resistance to partial NFS1 inhibition, exhibiting restored insulin function and metabolic flexibility, and attenuated loss of adiposity.

## Discussion

Here, we show that the main acute effects of acute *in vivo* inhibition of ISC biosynthesis are an unexpected shift to lipid utilization and rapid loss of adipose tissue without observable changes to activity or feeding. While over 60 proteins need one or more ISCs for their function or stability, the degree to which any given ISC protein is affected is likely protein and context-dependent. Oxygen level is a key determinant of ISC protein stability and *in vivo* models provide the opportunity to study ISC biosynthesis inhibition in physiological oxygen tension.^23,24^ While atmospheric oxygen tension is 21%, physioxia in internal organs varies from 14% in lung alveoli, ∼5.4% in liver^40,41^, ∼3.8% in skeletal muscle^41^, and 4.7-8.9% in adipose tissue.^42^ Moreover, tissue- and cell-type specific differences in ISC biosynthesis flux and ISC protein stability may also underlie differences in ISC protein levels observed in specific tissues.

We find that specific tissues differentially activate stress responses to cope with ISC deficiency. While ISCs play an important role in iron sensing, only the liver showed activation of the iron starvation. response upon inhibition of ISC biosynthesis, similar to the IRP-mediated response described *in vitro.*^7^ This differential activation may be related to the liver’s role as the main iron-sensing organ in the body. Interestingly, muscle and heart exhibit reduced Nfs1 knockdown efficiency despite these tissues exhibiting adequate knockdown efficiency in other similar models.^43^ These data could indicate a mechanism for ISC sensing and compensation in these tissues. Indeed, microarray analysis in muscle biopsies from patients with myopathy caused by mis-splicing of ISCU mRNA in skeletal muscle shows significantly increased NFS1 expression.^44^ We observed upregulation of Atf5, Nrf2, and Hmox1 exclusively in muscle upon ISC inhibition, indicating that this tissue could be hypersensitive to oxidative stress, and providing rationale for regulating ISC biosynthesis via Nfs1 activation.

Many of the transcriptional stress responses and metabolic phenotypes observed upon ISC inhibition in mice are similar to other mouse models of reduced mitochondrial function, reminiscent of Nfs1 patients that present with mitochondrial disease-like phenotypes at birth. Similar to mice with mitochondrial OXPHOS defects, we observe Atf4, Atf5, and PGC-1a upregulation and elevated plasma lactate. However, we observed several phenotypes of ISC synthesis inhibition that are not shared by models of defective OXPHOS. Surprisingly, the mitochondrial stress response induced by ISC synthesis inhibition is not accompanied by upregulation and secretion of the metabokines GDF15 and FGF21, which mediate an adaptive response to mitochondrial dysfunction.^32,34^ Indeed, while similarly activating the ISR, mtDNA Deletor mice upregulate FGF21 and shift to glucose as the major metabolic fuel, demonstrated by increased RER.^32^ Also, in patients with OXPHOS defects, elevated plasma lactate is often accompanied by elevated plasma insulin.^45^ These differences are likely related to the important role of ISCs in proteins beyond the ETC, including insulin secretion and mRNA transcription.

Thus, the main difference in mice with ISC dysfunction compared to defective OXPHOS alone may be that besides reduced mitochondrial function, mice with ISC dysfunction are unable to shift to glucose as their main fuel. In young mice, Nfs1 inhibition causes defects in insulin production, likely via reduced CDKAL1 levels, leading to major metabolic inflexibility. Interestingly, insulin can induce FGF21 expression in human skeletal muscle and increase circulating FGF21 levels^46^ and muscle FGF21 expression is associated with hyperinsulinemia.^47^ These findings might explain the unchanged Fgf21 expression in our mice despite activation of the ISR. Indeed, in muscle biopsies from patients with myopathy caused by ISCU mis-splicing, ISC proteins are unaffected in the pancreas, and a 25-fold induction of FGF21 mRNA is observed.^44^

There is an increasingly appreciated connection between iron metabolism and insulin production. A high frequency of diabetes is observed in patients with Friedreich’s Ataxia (20-50%) and iron overload disorders such as hereditary hemochromatosis (30–60%), phenotypes related to both insulin resistance and destruction of pancreatic β-cells.^48–51^ CDKAL1 is one of the most recently discovered ISC proteins, requiring two clusters^52^, and we show that it is reduced upon acute Nfs1 inhibition. Cdkal1 also plays a role in decreased insulin production in mice lacking iron-regulatory protein 2 (Irp2), which show decreased ISC biosynthesis and CDKAL1 levels in pancreatic islets.^53^ In humans, SNPs in CDKAL1 have been associated with low production of insulin and diabetes.^54–56^ Interestingly, the associations between CDKAL1 SNPs and diabetes are more pronounced in early-onset diabetes and are not found in late onset diabetes.^57,58^ Additionally, CDKAL1 risk alleles are associated with increased glucose levels from birth to 5 years of age.^59^ This age-related association between CDKAL1 and diabetes is in line with the differential metabolic effects of aged versus young mice upon ISC synthesis inhibition.

Ultimately, these age-dependent effects of *in vivo* ISC synthesis inhibition shed light on physiological interplay between iron metabolism and metabolic flexibility. This model and our findings might aid in better understanding the metabolic consequences in patients with ISC disturbances.

## Acknowledgements

We thank NYU Langone’s Experimental Pathology Research Laboratory (RRID: SCR_017928) who provided histologic processing and imaging of tissues, supported by the Cancer Center Support Grant (NIH/NCI 5 P30CA16087). We thank NYU Langone’s Genome Technology Center (GTC) (RRID: SCR_017929), Applied Bioinformatics Laboratory (RRID:SCR_019178) and Rodent Genetic Engineering Core (RRID:SCR_017925), supported by supported by the Cancer Center Support Grant (P30CA016087). This work has used computing resources at the High Performance Computing Core (HPC) of the Medical Center Information Technology group at the NYU Langone Medical Center. We thank NYU Langone’s Rodent Behavior Laboratory (RRID: SCR_017942) for providing metabolic cage experiments. We thank the Stable Isotope and Metabolomics Core Facility of the Diabetes Research and Training Center (DRTC) of the Albert Einstein College of Medicine, for performing metabolomics and lipidomics, supported by NIH/NCI grant (P60DK020541). We thank the Human Islet and Adenovirus Core of the Einstein-Mount Sinai Diabetes Research Center (P30 DK020541) for demonstrating how to perform islet isolations. MM is supported by an EMBO Postdoctoral Fellowship (ALTF 1104-2021) and an HFSP Long-Term Fellowship (881965). Part of this work was supported by Novo Nordisk.

## Author contributions

MM and RP conceived and designed the project. MM, HM, SWA, LA and DZ performed experiments. MM, HM, CR, CYP, AG, AMS and RP analyzed the data. MM and RP wrote the manuscript with contributions from all other authors.

## Supplemental information

Document S1. Figures S1–S4

Table S1. Excel file containing Transcriptomics data, related to Figure 1.

Table S2. Excel file containing Metabolomics data, related to Figure 3.

Table S3. Excel file containing Lipidomics data, related to Figure S3.

## Methods

### Doxycycline inducible shNfs1 mouse model description

Transgenic mice were generated by Mirimus as followed: Two transgenes were introduced into D34 embryonic stem (ES) cells: 1) a Tet-inducible shRNA targeting Nfs1 coupled with turboGFP (TRE-tGFP-shNfs1) and 2) a floxed reverse tetracycline-controlled transactivator (rtTA3) coupled with mKate2 (CAG-lox-stop-lox-rtTA3-IRES-mKate2). The shNfs1 transgene was knocked in downstream of Col1a1 on chromosome 11, and the rtTA3 transgene was knocked in at endogenous Rosa26 loci on chromosome 6. Transient expression of Cre recombinase in ES cells resulted in the excision of the lox-stop-lox cassette, enabling expression of rtTA3 and mKate2. ES cells were injected into tetraploid blastocysts to create a transgenic mouse on a mixed background that expresses the rtTA3 protein in all tissues and activates the expression of shNfs1 upon adding doxycycline. Control mice for all experiments harbored TRE-tGFP-shNfs1 but did not express rtTA3. As such, treatment of the control mice with doxycycline did not elicit shNfs1 expression or NFS1 inhibition. A separate cohort of mice in which the lox-stop-lox cassette is intact have been backcrossed to a C57BL/6 background and are available for studying tissue-specific effects.

### Mouse maintenance conditions

Mice were fed a standard laboratory rodent diet throughout their life cycle, and housed in a room with a 12 hour light-dark cycle. For the high fat diet experiment, mice were fed either a rodent diet with 60 kcal% fat (D12492, Research Diets Inc) or a rodent diet with 10 kcal% fat (matching sucrose to D12492, Research Diets Inc). The amount of doxycycline used to induce effects was titrated down to 50 ug/mL via drinking water (8 mg/kg/day), >20-fold less than the amount used to induce shRNAs in most tumorigenesis models or cause microbiome dysbiosis (200-320 mg/kg/day).^23,60^ Both control mice and mice in which Nfs1 can be inhibited receive doxycycline treatment. All experiments included both female and male mice, except for the high fat diet experiment, which included only male mice.

### Western blotting

For protein isolation, snap frozen tissues were homogenized with a homogenizing pestle in RIPA lysis buffer (50 mM tris (pH 7.4), 150 mM NaCl, 1% NP-40, 0.1% sodium deoxycholate, 0.1% SDS, and 2 mM EDTA) with a protease inhibitor cocktail (Sigma-Aldrich) on ice. Lysates were sonicated at an amplitude of 25% in 15 second pulses (15s on, 15s off) on 4C for a total of 1.5 minutes using Model 120 Sonic Dismembrator (Fisherbrand). Lysates were centrifuged at 4°C at 20,000g for 10 min and protein concentrations of the supernatants were determined by Pierce BCA Protein Assay Kit (Pierce). Sample loading buffer was added to 7 μg of protein per lysate, boiled for 5 min, and loaded into Bolt 4 to 12% bis-tris polyacrylamide gels (Fisher). Gels were run on 100V for 1.5 hours, and transferred to a PVDF Transfer Membrane (Millipore) at 60V for 3.5 hours. Depending on the primary antibody, membranes were either blocked with 5% bovine serum albumin (BSA), or 5% milk (for GAPDH antibody), and membranes were incubated with primary antibodies at 4C overnight. Membranes were washed with TBS-T (tris buffered saline, 0.1% Tween-20), incubated with secondary antibody for 1 hour at room temperature, washed with TBS-T, and developed using ECL substrate (Thermo Fisher Scientific) with autoradiography film (WorldWide) in dark room. Primary antibodies used were NFS1, B-ACTIN, α-Tubulin, SDHB, FECH, CDKAL1, GAPDH, LIAS. Bands were quantified using ImageJ.

### Blood Counts

Blood cell counts were measured using the Element HT5 (Heska). Mice were anesthetized in an induction chamber using isoflurane. Blood was collected from the submandibular facial vein using a 5mm lancet. 100 μl of blood was collected into a K_2_EDTA-coated tube and gently inverted to prevent clotting and to prevent excessive RBC lysis. Measurements were taken within 30 minutes of blood collection.

### Bone marrow isolation and transplant

Bone marrow was isolated from five control mice. Following euthanasia, mice were dissected and both femurs and tibias were removed and collected in a 50mL Falcon tube containing ice cold HBSS. The ends of each bone were cut using bone cutting scissors and the bones were placed into a 1.75mL Eppendorf tube containing 500uL ice-cold HBSS. The tubes were then centrifuged at 10,000G for 30 seconds at 4C. The bone marrow was pooled into a 50mL tube after which the suspension was passed through a 50 um cell strainer. After centrifuging at 350g for 10 min at 4C, the supernatant was aspirated in the pellet was resuspended in fresh HBSS. Cells were counted and appropriate volume was resuspended in 200uL of ice cold HBSS and kept on ice until transplantation. Nfs1 KD and control mice were irradiated with 9gy of radiation using a MultiRad350 irradiator (Precision X-Ray). Then mice were anesthetized and the 200uL bone marrow suspension was injected into the retro-orbital vein using a 25g needle. A topical eye antibiotic ointment was applied and the mice were then returned to the cage and observed during recovery. For the following two weeks, mice were kept on antibiotics to prevent infection, after which mice were treated with doxycycline supplemented water (50 ug/mL) to initiate NFS1 knockdown.

### Histopathology and islet density analysis

Mice tissues were fixed in PFA for 24 hour at 4 °C. Five-micron sections of paraffin embedded tissue were deparaffinized and stained with hematoxylin and eosin (Η&E) or periodic acid-Schiff (PAS). Pancreatic islets and nuclei were identified and segmented by size and shape using CellProfiler, and their respective areas were calculated.

### RNA isolation

For mRNA isolation, snap frozen tissues were homogenized with a homogenizing pestle in TRIzol (Invitrogen) and isolation was continued according to manufacturer’s protocol. For RNaseq, contaminating genomic DNA was removed using RNase-Free DNase (QIAGEN) and samples were cleaned up with the RNeasy Mini Elute Cleanup Kit (QIAGEN).

### Library Preparation

RNA extractions were quantified using RNA Nano Chips (Agilent) on an Agilent 2100 BioAnalyzer. RNA-Seq library preps were constructed using the Illumina TruSeq Stranded mRNA Library Prep kit (Illumina) using 100ng of total RNA as input, amplified by 14 cycles of PCR. Final libraries were visualized using High Sensitivity DNA ScreenTape (Agilent) on the Agilent TapeStation 2200 instrument. Quant-It (Invitrogen) was used for final concentration determination and libraries were pooled equimolar. The pool was sequenced paired-end 50 cycles on a single lane of an Illumina NovaSeq6000 S1 100 Cycle flowcell-v1.5 with 2% PhiX spike-in.

### Read Mapping, Statistical Analyses, and Data Visualization

Per-read per-sample FASTQ files were generated using the bcl2fastq2 Conversion software (v.2.20) to convert per-cycle BCL base call files outputted by the sequencing instrument into the FASTQ format. The alignment program, STAR (v2.7.3a), was used for mapping reads of 23 samples to the mouse reference genome mm10 and the application Fastq Screen (v0.13.0) was utilized to check for contaminants. The software, featureCounts (Subread package v1.6.3), was used to generate matrices of read counts for annotated genomic features. For differential gene statistical comparisons between groups of samples contrasted by Nfs1 and Control conditions across liver, muscle, and white adipose tissues, the DESeq2 package (R v4.1.2) in the R statistical programming environment was utilized. In R, PCA plots were generated with the prcomp() function of the stats package along with the ggplot2 package. The raw sequencing data were deposited to the NCBI Sequence Read Archive (accession number SRA: PRJNA1129102).

### Metabolic Cages and DEXA scan

To measure respiratory exchange ratio (RER), four mice per condition were single housed in metabolic cages (TSE PhenoMaster), in which activity and food/water intake was simultaneously reported. Mice were acclimatized to the metabolic cages for at least two days and analyzed for two or more days. After metabolic cage analysis, body composition was measured in anesthetized mice by dual-energy X-ray absorptiometry (DEXA) scans, according to the manufacturer’s instructions (Scintica).

### Plasma Lipidomics

A volume of 30 μl of plasma was used for the assay. Samples were extracted in 9 volumes of ethanol with internal standards. The samples were centrifuged, and the supernatant were transferred to another vial and dried under gentle nitrogen flow. Dried samples were then dissolved in 100ul ethanol for injection. Samples were analyzed on the ABsciex 6500+, and a pooled quality control (QC) sample was included in each run. This QC sample was injected six times for coefficient of variation (CV) calculation for data quality control. A total of 482 lipids were detected with CV less than 30% in QC samples.

### Plasma Metabolomics

For plasma samples, a volume of 30 μl of plasma was used for the assay. The samples were extracted with 120 μl of ethanol with internal standards. The samples were centrifuged, and supernatant was transferred to another vial for injection. Samples were analyzed with ABsciex 6500+, and a pooled quality control (QC) sample was included in each run. This QC sample was injected six times for coefficient of variation (CV) calculation for data quality control. A total of 296 small metabolites were detected with CV less than 30% in QC samples.

### Lipidomics and Metabolomics analysis

Normalized data sets were imported into SIMCA-p software (Umeå, Sweden) for multivariate analysis. Unsupervised principal component analysis (PCA) and supervised partial least square-discriminant analysis (PLS-DA) were performed to analyze separation among groups (fast/refed, control/shNfs1). Variable importance in the projection (VIP) from the PLS-DA analysis was calculated. The average, fold change, and p values using the student’s t test were calculated.

### Glucose metabolism

Fasted (6 hours) glucose was measured using a glucometer (Care Touch) from blood taken by snipping the tip of the animal’s tail. Circulating plasma hormones were determined using the Mouse/Rat Metabolic Hormone Discovery Assay® Array (Eve Technologies; Canada). Blood was withdrawn from mice that were fasted for 4 hours by puncturing the sub-mandibular vein into an EDTA-coated tube, and 55μL of Plasma was shipped to Eve Technologies on dry ice.

For the oral glucose tolerance test (OGTT) mice were fasted during the daytime for 5 hours prior to the OGTT. A stock solution of 50% dextrose (Durvet) was diluted to 12.5% in 0.9% saline, and the mice were gavaged with a dose of 2g dextrose/kg of mouse weight. For every time point, the first drop of blood was wiped off and a reading was taken from the following drops, after which the tail was wiped with gauze. Subsequent readings were taken at: 0, 15, 30, 45, 60, 90, and 120 minutes. To minimize the impact of stress on blood glucose, the mice were allowed to roam freely in their cages as their blood was taken. Using the first blood glucose measurement as the baseline, AUCs were calculated for each mouse using GraphPad (Prism) and significance was calculated with student’s t-test.

Similar to the OGTT, mice were fasted during the daytime for 5 hours prior to the insulin tolerance test (ITT). For the ITT, a stock of Humulin-100 (100 IU/mL) was diluted to 0.15 IU/mL in 0.9% saline. 0.75 U of insulin/kg of mouse weight was injected via intraperitoneal injection using SafetlyGlide 31G insulin needles (BD). Blood glucose measurements were performed as described for the OGTT. In case of hypoglycemia (blood glucose dropping below 50 g/dL), mice were injected with 300uL of a 10% dextrose solution (Durvet) and animals were censored.

### Islet isolation

Islet isolation was performed as described^61^ originated from.^62^ In brief, mouse islets were isolated by clamping the bile duct after which Hank’s buffered saline solution containing collagenase P was injected through the pancreatic duct at the ampulla of Vater. After successful injection confirmed by fully inflated pancreata, the pancreata were collected and digested at 37°C, and separated by density gradient in Histopaque (Sigma). Islets were washed with Hank’s buffered saline solution and snap frozen and stored at −80C until further processing.

