## Supplemental Figures for "Acute inhibition of iron-sulfur cluster biosynthesis disrupts metabolic flexibility in mice"

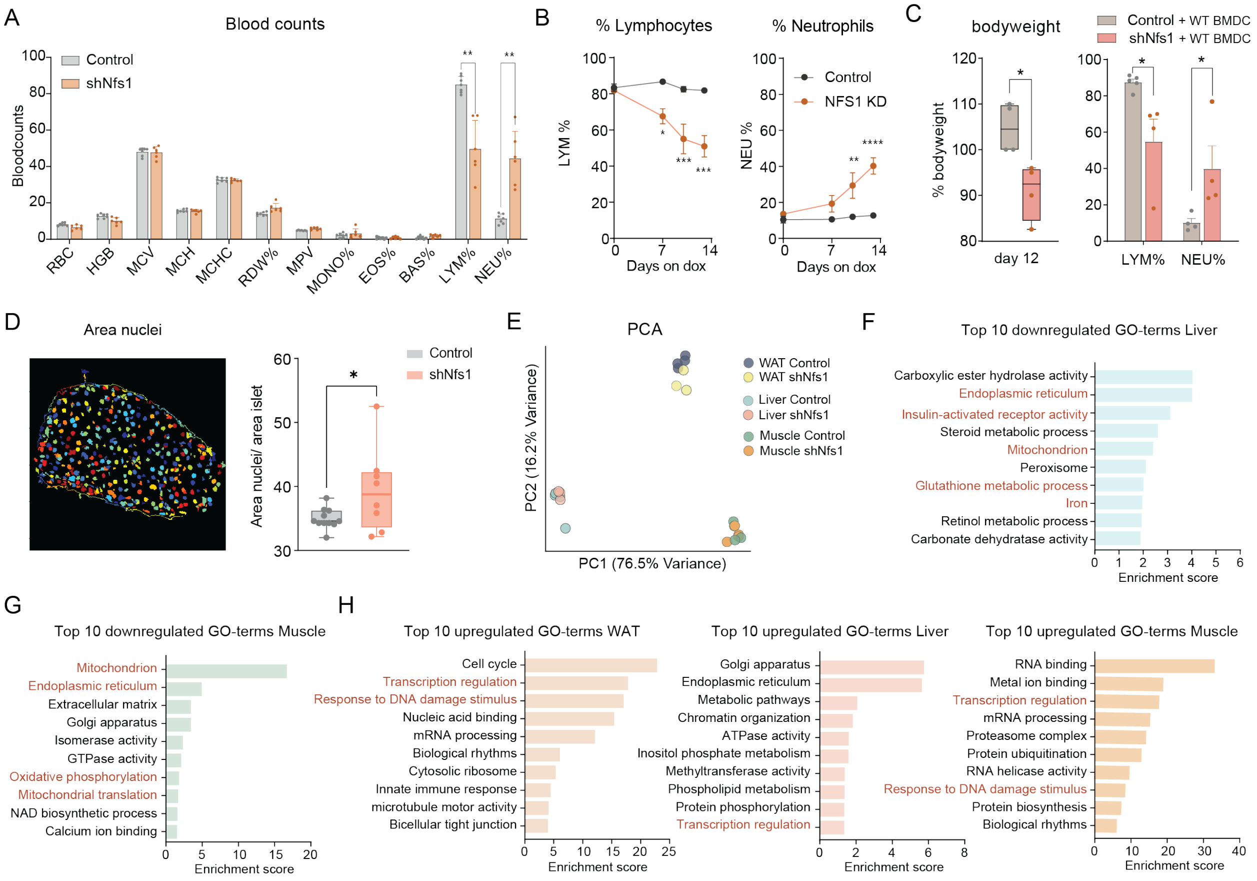


**Figure S1. related to Figure 1.**

A) Blood counts 12 days after dox addition, n=7 for control and n=6 for Nfs1-suppressed mice.

B) LYM% and NEU% at indicated days after dox addition, n=4.

C) Bodyweight (%), LYM% and NEU% 12 days after dox addition in bone marrow-transplanted control and Nfs1-suppressed mice, n=4.

D) Area nucei/ area of the islets from H&E stained pancreas slides; depiction of area selection is depicted on left side, n=11 for control and n=8 for Nfs1-suppressed mice.

E) PCA analysis on RNAseq data.

F) The top 10 enriched GO-terms downregulated in Liver and G) Muscle.

H) The top 10 enriched GO-terms upregulated in eWAT, Liver and Muscle.

Data are presented as mean ± SEM; comparisons were made using two-way analysis or student's t-test (C, body weight% and D). $^*p < 0.05$, $^{**}p < 0.01$, $^{***}p < 0.001$, $^{****}p < 0.0001$.

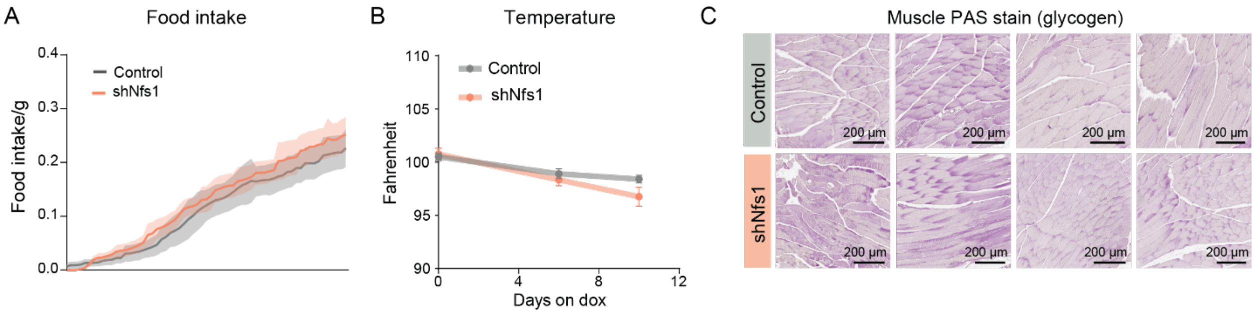


**Figure S2, related to Figure 2.**

A) Caloric intake per g of bodyweight measured in shNfs1 and control mice, with N=4 per group.

B) Body temperature measured at room temperature during different time points between 0 and 10 days of doxycycline.

C) Periodic acid-Schiff reagent (PAS) demonstrates the amount of glucose stored as glycogen in Muscle of control and Nfs1 KD mice after 9 days of doxycycline.

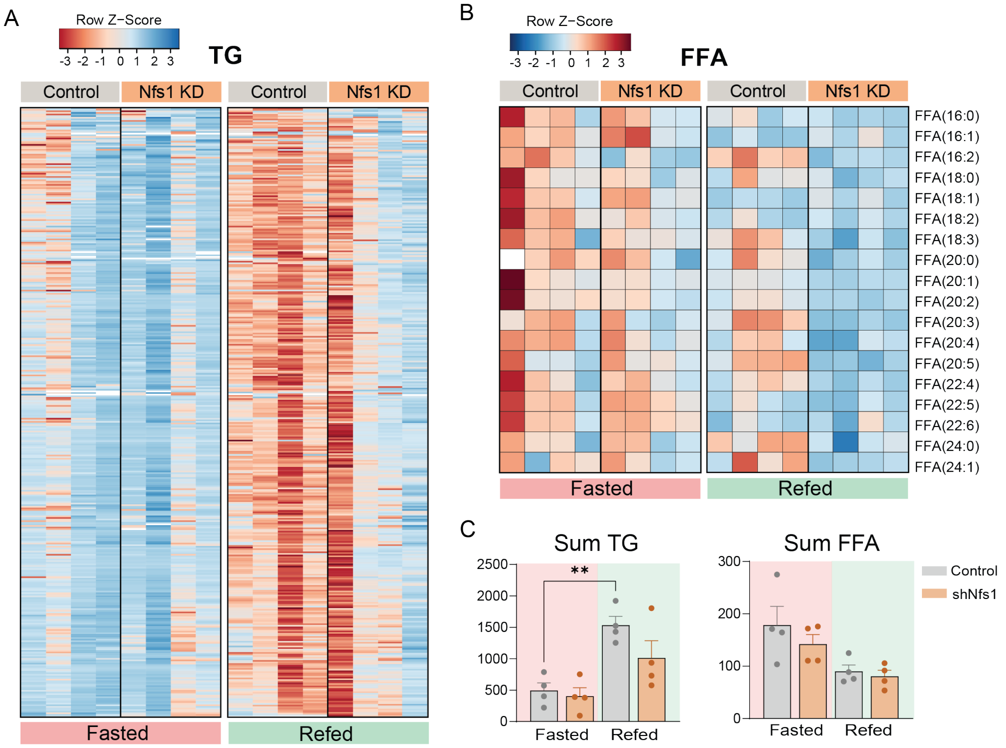


**Figure S3, related to Figure 3.**

A) Heat map of all triglyceride (TG) species detected in plasma during fasted and fed state and upon Nfs1 inhibition vs control mice, measured by lipidomics.

B) Heat map of all free fatty acid (FFA) species detected in plasma during fasted and fed state and upon Nfs1 inhibition vs control mice, measured by lipidomics.

C) Sum of all TGs and sum of all FFS detected in plasma during fasted and fed state and upon Nfs1 inhibition.

Data are presented as mean ± SEM; comparisons were made using two-way analysis of ANOVA, **$p < 0.01$, with N=4 for all groups.

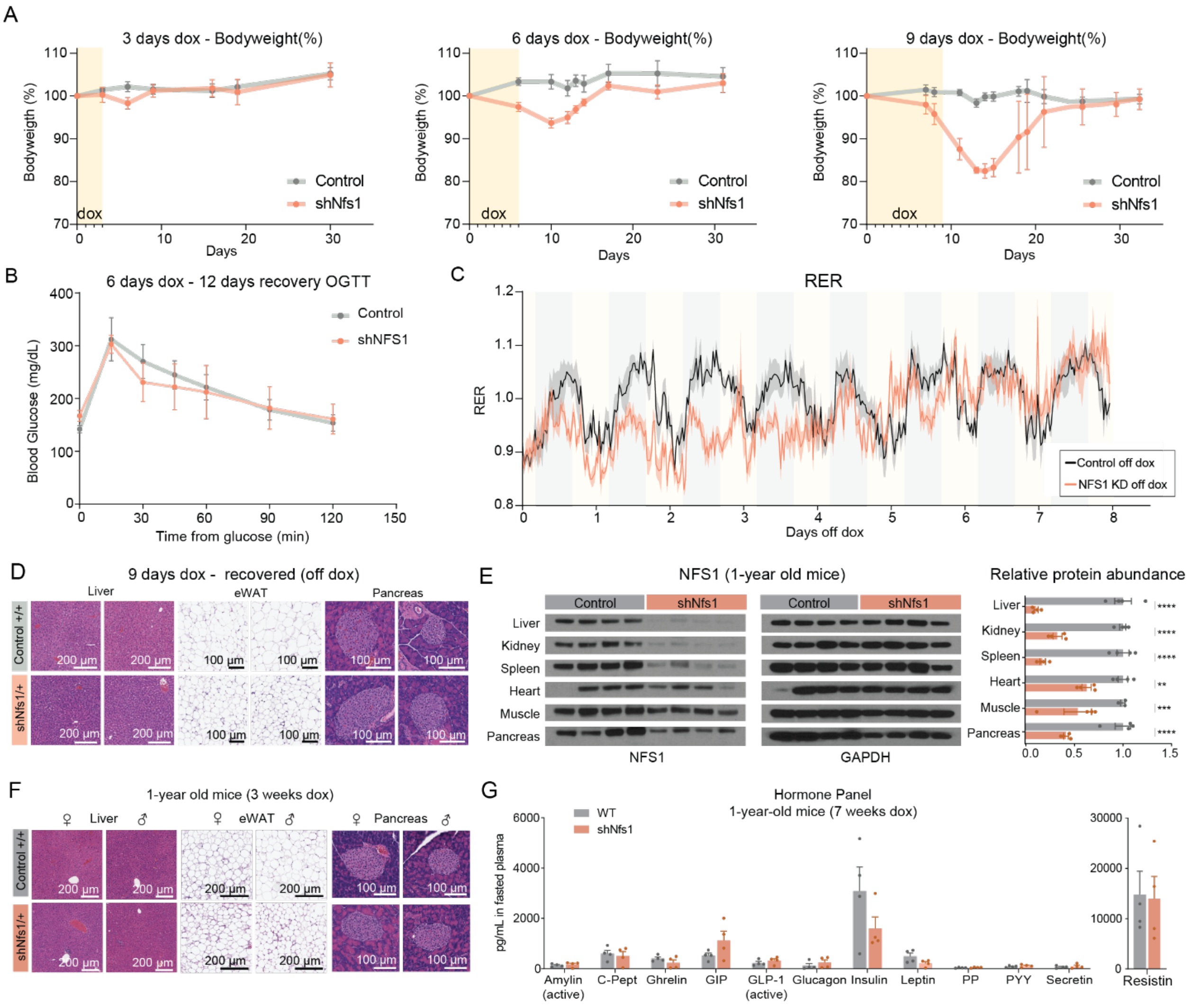


**Figure S4, related to figure 4.**

A) Bodyweight in control and Nfs1 KD mice treated with 3, 6, or 9 days of doxycycline and withdrawal.

B) Oral Glucose tolerance test (OGTT) for shNfs1 and control mice treated with 6 days of dox which have recovered for 12 days. Blood glucose concentrations were determined at indicated times.

C) Respiratory exchange ratio (RER, $VCO_2/VO_2$) measured over an 8-day period after 6 days of doxycycline exposure.

D) Hematoxylin and eosin stain (H&E) shows histology of liver, eWAT, and pancreas of control and Nfs1-inhibited mice treated with 9 days of doxycycline and recovery upon withdrawal.

E) Western blot analysis of NFS1 protein levels in extracts of livers, kidneys, spleens, muscles, hearts, and pancreas from control and Nfs1 KD mice. Quantitation of protein levels normalized to loading control shown at right.

F) Hematoxylin and eosin stain (H&E) shows histology of liver, eWAT, and pancreas of 1-year old control and Nfs1-inhibited mice after 3 weeks of dox.

G) Metabolic hormone panel from 4 hour fasted plasma from 1-year old control and Nfs1-inhibited mice after 7 weeks of dox, with n=4 for both groups.
